# Active Sampling and Sex Differences in Perceptual Decision Making in Rats

**DOI:** 10.1101/2025.10.25.684539

**Authors:** Jensen A Palmer, Kevin Chavez Lopez, Mark Laubach

## Abstract

Decisions are often modeled as a sequential process in which evidence accumulates until it reaches a threshold, triggering a response. Studies in freely moving animals raise questions about how ongoing behavior, not just stimulus properties, shapes this process. We trained rats of both sexes on a visual detection task with three luminance levels, each associated with the same reward outcome. Rats controlled cue duration through sustained head entries into a center port, yielding a measurable index of active sampling. Females consistently sampled longer than males. Sampling durations were shorter on error than correct trials, and reaction times were longer on error trials. We used drift diffusion models to relate these behaviors to the decision process. Luminance selectively affected the rate of evidence accumulation, with drift rate increasing monotonically across low, mid, and high luminance levels. Active sampling time was associated with the decision threshold, with longer sampling predicting higher thresholds in both sexes. The relationship between sampling time and drift rate differed by sex. Females showed a negative association between sampling duration and drift rate that was absent in males. These findings suggest that cue properties and active sampling make separable contributions to decision making, with a negative association between sampling duration and drift rate evident in females but not males.

---

The standard framework for perceptual decision making treats perception, decision, and action as sequential stages. Sensory information is received and processed, a decision is formed, and a motor response follows. Models like the Drift Diffusion Model (DDM) are built on this framework (Ratcliff, 1978) and have been widely applied to understand the computational basis of decision making in humans and animals. These models assume that noisy sensory evidence accumulates over time until it reaches a threshold, triggering a response. Within this framework, the quality of the sensory signal determines the rate of evidence accumulation, and the threshold determines how much evidence is required before a decision is made.

This framework emerged from human experiments in the 1970s. Similar paradigms remain in use today (e.g., Johansson and Ulrich, 2025). Participants are presented with stimuli and make discrete motor responses, such as eye or hand movements, to report their decision. In contrast, experiments with freely moving animals are more complex (Carandini and Churchland, 2013). Animals must be trained to move to specific locations in a behavioral arena, producing task stimuli through actions like head entries into response ports. They report their choice by moving to another location. They then receive food or fluid reward, often at a separate location. These differences imply alternative interpretations of DDM parameters for studies in freely moving animals.

An alternative to the classic sequential framework comes from ecological psychology (Gibson, 1966), which proposes that perception is an active process and that decisions emerge from an ongoing cycle of actions, perceptions, and further actions. More recent formulations of this perspective include the affordance competition hypothesis (Cisek, 2007), the embodied choice framework (Lepora and Pezzulo, 2015), and active inference frameworks (Linson et al., 2018). Within sequential sampling models, this view implies that animals actively sample their environment. Sampling behavior itself may be part of the evidence accumulation process rather than a precursor to it. This possibility motivates two questions addressed in the present study: whether active sampling contributes to decision making in freely moving rats and which computational parameters it influences.

Our group has used a series of tasks to examine visually guided decisions in rats. White et al. (2024) examined how rats initially learn to make choices between visual stimuli paired with different reward values. Palmer et al. (2024) found that reversible inactivation of the rodent prelimbic cortex selectively increased the decision threshold without affecting drift rate, and identified sex differences in the rate at which decision thresholds changed across sessions of choice learning. Male rats showed greater reductions in threshold over time, consistent with higher choice impulsivity. Palmer et al. (2025) addressed a design limitation of these earlier studies by introducing a third stimulus of intermediate luminance to separate the effects of visual salience from reward value. That study found that reducing the perceptual discriminability between options reduced drift rate and starting point bias without affecting threshold. These findings demonstrate a dissociation between the computational effects of sensory salience and prefrontal function. However, in all three of these studies luminance was paired with reward value, making it difficult to isolate the contribution of luminance itself to the decision process.

The present study addresses this limitation directly. We trained rats on a visual stimulus detection task in which three luminance levels were all associated with the same reward outcome, removing the value confound entirely. We also required rats to maintain sustained head entries in the center port to produce the task stimuli, making the duration of active cue sampling a behavioral variable we could measure and analyze. This design allowed us to ask two questions. First, does luminance affect DDM parameters in the same way when reward value is held constant? Second, does the duration of active sampling contribute to decision making, and if so, which parameters does it influence? We found that luminance selectively affected drift rate, replicating the findings of Palmer et al. (2025) in a pure detection context. Active sampling time was associated with the decision threshold, with longer sampling predicting higher thresholds in both sexes. We also identified a sex difference in the relationship between sampling time and drift rate that is consistent with the impulsivity-related sex differences observed in our previous work.

## Methods

All procedures were approved by the Animal Care and Use Committee at American University (Washington, DC). Procedures conformed to the standards of the National Institutes of Health Guide for the Care and Use of Laboratory Animals. All efforts were taken to minimize the number of animals used and to reduce pain and suffering.

### Animals

Rats were kept individually on a 12-hour light and 12-hour dark cycle. During training and testing, they had controlled access to food (12-16 grams) to keep their body weights at about 90% of what they would weigh with unlimited food. Two groups of rats were used in these experiments. The first group had seven female (200-250g) and seven male (300-400g) Long Evans rats, obtained from Charles River. We trained this group using our initial procedure, where the task stimuli only stayed active as long as the rats kept responding in the center port. However, only four females and five males progressed during training without developing spatial biases. Some of the rats who eventually learned to perform the task without spatial bias required sessions with unbalanced presentations of left and right cues to overcome spatial bias. Results from the subset of rats who eventually did not show frank spatial bias are reported below. We modified the task design and trained a second cohort of rats, eleven females and eleven males from the same vendor and with the same weight ranges as above. These rats were trained with longer cue durations, as described below, and all of them met the training criteria without showing spatial biases.

### Behavioral apparatus

Animals were trained in sound-attenuating behavioral boxes (Med Associates) that had a single, horizontally placed spout mounted to a lickometer 6.5 cm from the floor with a single white LED placed 4 cm above the spout. Solution lines were connected to 60cc syringes and solution was made available to animals by lick-triggered, single speed pumps (PHM-100; Med Associates) which drove syringe plungers. Each lick activated a pump which delivered roughly 30 μL per 0.5 second activation. On the wall opposite the spout, three 3D-printed nosepoke ports were aligned 5 cm from the floor and 4 cm apart, and contained Adafruit 3-mm IR Break Beam sensors. A Pure Green (530 nm) 1.2” 8×8 LED matrix (Adafruit) was placed 2.5 cm above the center of each of the three nose poke ports outside of the box for visual stimulus presentation. LED matrices are controlled using Adafruit_GFX and Adafruit_LEDBackpack libraries, and microcontroller software is provided in a previous publication (Swanson et al., 2021).

### Behavioral task

The sequence of events in the behavioral task is shown in Figure 1A. Rats were initially trained to lick on a recessed spout within a reward port to receive 8% wt/vol liquid sucrose (30 µl). A white LED above the reward port indicated when rewards were available. In subsequent sessions, animals were trained using the method of successive approximations to respond in nose poke ports on the opposite wall of the behavioral arena to produce rewards at the reward port. A 4x4 square of green LEDs displayed over the center of the three ports signaled available trials. When a trial began, one of three cues was presented randomly on either the left or right side. The lateralized cues stayed on until rats entered a side port. To trigger a reward at the reward port, rats had to enter the signaled side port.

**Figure 1.**
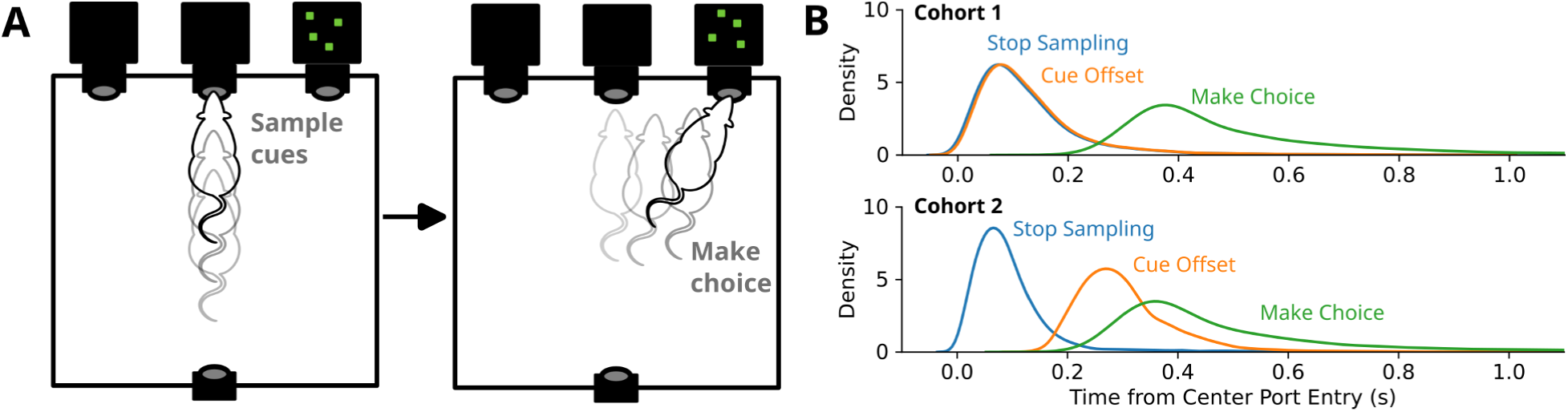
Task design. This figure describes the behavioral design and the timing of the task events including the sampling period. **A.** In decision-making tasks with freely moving rodents, animals must enter a center port to trigger a stimulus and move to a lateral port to report their choice. This setup requires active head and body positioning, engaging a wide range of body parts to complete the task. In the task used in this study, rats produced cues by making sustained head entries into a center port, called sampling behavior. Then, they reported the location of the cues by moving to a lateralized port. The task used dynamic visual stimuli presented with 8x8 grids of green LEDs located above the response ports. **B.** Relative timing of sampling, cue duration, and response time for the two cohorts of rats. In the first cohort, sampling controlled cue duration and many rats struggled with spatial bias early in training. In the second cohort, cue duration was extended based on the minimal response time and spatial bias was minimal.

Correct responses (detections or hits) to any cue yielded access to 8% wt/vol sucrose reward at the reward spout. Responses to non-illuminated ports were considered as errors (misses) and were unrewarded. Rats were required to travel to the reward port and contact the spout after errors. This maintained a common sequence of behavioral events across trials. Choices (correct or incorrect) that took longer than 5 seconds following trial initiation were counted as errors of omission and were unrewarded. On valid trials, animals had to collect their reward within 5 seconds following responses to receive fluid. Errors of omission and trials with very delayed reward retrieval were rare (less than 5% of trials). Animals were trained in this task until they demonstrated consistent behavior (detection rates, sampling times, etc.) over three consecutive testing sessions, which comprised the data set for all subsequent analyses.

Three levels of luminance were used: high-luminance (16 LEDs), mid-luminance (four LEDs), or low-luminance (one LED). The location of illuminated LEDs within the 8x8 grid changed every millisecond, making for dynamic visual stimuli. The luminance of the cue on a given trial was chosen randomly. Luminance levels, measured using a S170C photodiode sensor and ThorLabs PM100D photometer, with static stimuli were 14 µW (high), 3.8 µW (mid), and 1.1 µW (low). (It was not possible to obtain accurate, stable measures of luminance using the dynamic visual stimuli given their flux rate of around 1 ms.) The luminance levels were selected to maintain detection rates above 90% for the high luminance cue in sessions early in training in which the cue stayed on until the rat entered one of the side ports.

After animals responded to all three cues with over 90% accuracy, the cue duration was reduced to the duration of the response in the center port. The active production of the cues by the rats’ sustained response in the center port was called "sampling". One cohort of rats was trained and tested using cues that were only available while the rats sampled the cues. Some rats in this cohort struggled with consistent performance, and developed spatial biases. To overcome this issue, we then tested an additional cohort of rats with temporally extended cues. We added the minimum response time from the previous testing session to the sampling time. If rats were able to respond with over 90% accuracy to all cues, we reduced the temporal padding of the cues to 75% of the previous session’s minimum response time. This resulted in longer stimulus times that were adjusted based on each rat’s performance. The relative timings of sampling, cue duration, and response time for the two cohorts of rats are shown in Figure 1B.

The behavioral measures of interest are referred to as “sampling time,” “cue duration,” “choice time,” “detection rate,” “reward retrieval,” and “inter-trial interval.” Sampling time, as previously stated, was the amount of time animals chose to sustain a response in the center port. Cue duration was the sampling time plus the minimal reaction time from the previous behavioral session. Choice time was measured as the time taken to respond in either the left or right side port after exiting the center port. Detection rate was the ratio of correct responses for each cue. Reward retrieval was measured as the time it takes from entering one of the side ports to the time of the first lick at the reward port. Inter-trial interval (ITI) was measured as the time from one trial initiation to the next.

### Statistical Analyses

Behavioral events were recorded through MED-PC and extracted through custom scripts written in Python (Anaconda distribution: https://www.continuum.io/). Statistical analyses were carried out using the pingouin package for Python (Vallat, 2018) and R (R Core Team, 2023) run in Jupyter notebooks (http://jupyter.org/) via rpy2. Mixed and repeated-measures ANOVAs were performed using Pingouin and the ezANOVA function in R. Within-subject effects included cue (high-, mid-, or low-luminance) and response (correct or error) and the between-subject effect was the rats’ sex. Logarithmic transformations were done where appropriate to account for skewed distributions. Reported statistics include p-values and F-statistics. Greenhouse-Geisser corrections and epsilon values are reported where instances of sphericity are violated. Where applicable, pairwise post hoc comparisons were performed with Tukey HSD adjustments. (R: “emmeans”, “pairs”).

For the Bayesian analyses reported below, we assessed our results using the mean estimates of all model parameters and the 94% Highest Density Interval (HDI 3% and 97%) of the posterior distributions of the parameters. For categorical factors (sex and cue luminance), instances where the 94% HDIs associated with a given level of the factor did not overlap the mean parameter estimate of another level of the factors were considered as significantly different. For regression using sampling time, we interpreted regression coefficients as significant if the HDIs did not overlap zero.

### Binary Choice Models

The Bambi package (Bayesian Model-Building Interface; Capretto et al., 2022) was used to model choice probabilities. Analyses were conducted using Python version 3.11.13, IPython version 9.3.0, and the following libraries: arviz 0.19.0, bambi 0.15.0, numpy 1.26.4, pymc 5.23.0, and pandas 2.3.0.

Hierarchical Bernoulli regression models were made for the binary choice data (coded as 1 for hits and 0 for misses). Model formulas were constructed using default priors to evaluate the influence of a categorical variable (cue luminance) and/or continuous variable (sampling time) on the probability of stimulus detection. Sampling time was robustly z scored using the median and median absolute deviation. The trained models provided full posterior distributions for each parameter. Models were run with four chains of 1000 tuning steps and 1000 draws. Target acceptance was increased to 0.95 from the default of 0.8 to improve convergence. Model fits were checked using the Gelman statistic (Gelman and Rubin, 1992), which was below 1.02 for all reported models, and using posterior predictive checks of the estimated choice probabilities. The autocorrelations and distributions of the parameters were visually assessed to further ensure convergence.

Following the fitting of each model, its performance and comparisons against alternative models was accomplished using the ArviZ library (Kumar et al., 2019). Model assessment was performed using the Pareto Smoothed Importance Sampling Leave-One-Out cross-validation (PSIS-LOO) (Vehtari et al., 2017). This technique estimates the out-of-sample predictive accuracy of a model. Pointwise LOO values were checked for all trials, ensuring the validity of our subsequent comparisons. Model comparison was then conducted using arviz.compare, which evaluates the expected log pointwise predictive density (ELPD) for each model. This allows for a comparison of models based on their out-of-sample predictive performance.

### Drift Diffusion Models

The HSSM package (version 0.2.5; https://doi.org/10.5281/zenodo.17247695; based on Fengler et al., 2026) was used to quantify effects of cue and sex on decision making. We estimated the four main DDM parameters, the drift rate, decision threshold, starting point bias, and non-decision time. Standard interpretations of these parameters are: Drift rate accounts for how quickly the rats integrate information about the cues. Threshold accounts for how much information is needed to trigger a choice. Starting point bias accounts for variability in the starting point of evidence accumulation. Non-decision time accounts for the time taken to process the cue and execute the choice.

Hierarchical HSSM models were fit that allowed for a single DDM parameter (drift rate, threshold, starting point bias, non-decision time) to vary freely across one of two factors (sex or cue luminance) or over the range of the sampling times. Other parameters were estimated globally. Models were run using the same Python environment described above for the binary choice models. Models were trained with 4 chains, with 1,000 tune and 1,000 draw iterations. Target acceptance was increased to 0.95 from the default of 0.8 to improve convergence. Models were fit using “choice time” as the measure of reaction time. Choice time was defined as the time from when the rats exited the center port to when they entered one of the side ports. Model comparisons used the same routines from the ArviZ package used for the binary choice models.

For all DDMs, convergence was validated based on the Gelman-Rubin statistic (Gelman and Rubin, 1992), which was below 1.02 for all parameters reported in this paper, with posterior predictive check that compared response times from the rats and the models, and by plotting the probability density functions for the observed and predicted response times for each rat. The autocorrelations and distributions of the parameters and predictions of the response latency distributions for each animal were visually assessed to further assess convergence.

## Results

### Behavioral Performance and Sex Differences in Sampling

Among the 9 rats from the initial cohort that completed training without persistent spatial bias, sampling duration differed by sex (F(1,7)=12.03, p=0.010), with females sampling longer than males. Sampling duration also differed by trial outcome (F(1,7)=12.93, p=0.0088), with longer sampling on hit than miss trials. The sex by outcome interaction was not significant (F(1,7)=1.00, p=0.350). By contrast, sampling duration did not vary systematically with cue luminance. Thus, longer sampling was associated with successful detection but not with the physical intensity of the cue. This pattern is consistent with the idea that accurate performance depended on sustained temporal engagement with the stimulus, rather than sampling duration being determined simply by cue luminance.

Mean sampling durations were brief, averaging approximately 184 ms in females and 77 ms in males. These durations are similar to those reported by Molano-Mazón et al. (2024), who also used sustained actions during stimulus presentation in a study of decision making by rats. Under this initial limited-sampling design, some rats developed persistent spatial response biases, identified using McNemar’s tests, and never achieved detection rates above 60% correct during training. These animals were not included in the analysis reported above. Overcoming these biases required extensive additional training that included blocks of trials with repeated spatial configurations. The resulting variation in training history, together with persistent spatial biases in some animals, raised concern that rats might not be using a common behavioral strategy. We therefore modified the task as described in the Methods and report results from a second cohort trained with the revised task below.

The second cohort of rats (n=22) was trained to perform the modified task, in which we added some time to the cue duration. The time added was based on each animal’s minimum response time (median of 196 ms, 95% CI: 158-350 ms), measured from the previous behavioral session. In this cohort, we found that the rats more quickly learned the task and did not demonstrate major spatial biases in performance. Female rats again sampled for longer than males, around 100 ms in the females and 75 ms in the males (p=8.793e-03, F(1,20)=8.428) (Figure 2B, left). The difference in sampling time was less dramatic with the use of the extended cues. Importantly, cue duration did not differ by sex (p=0.305, F(1,20)=1.107) (Figure 2B, right), allowing us to make direct comparisons between females and males in terms of other measures of performance.

**Figure 2.**
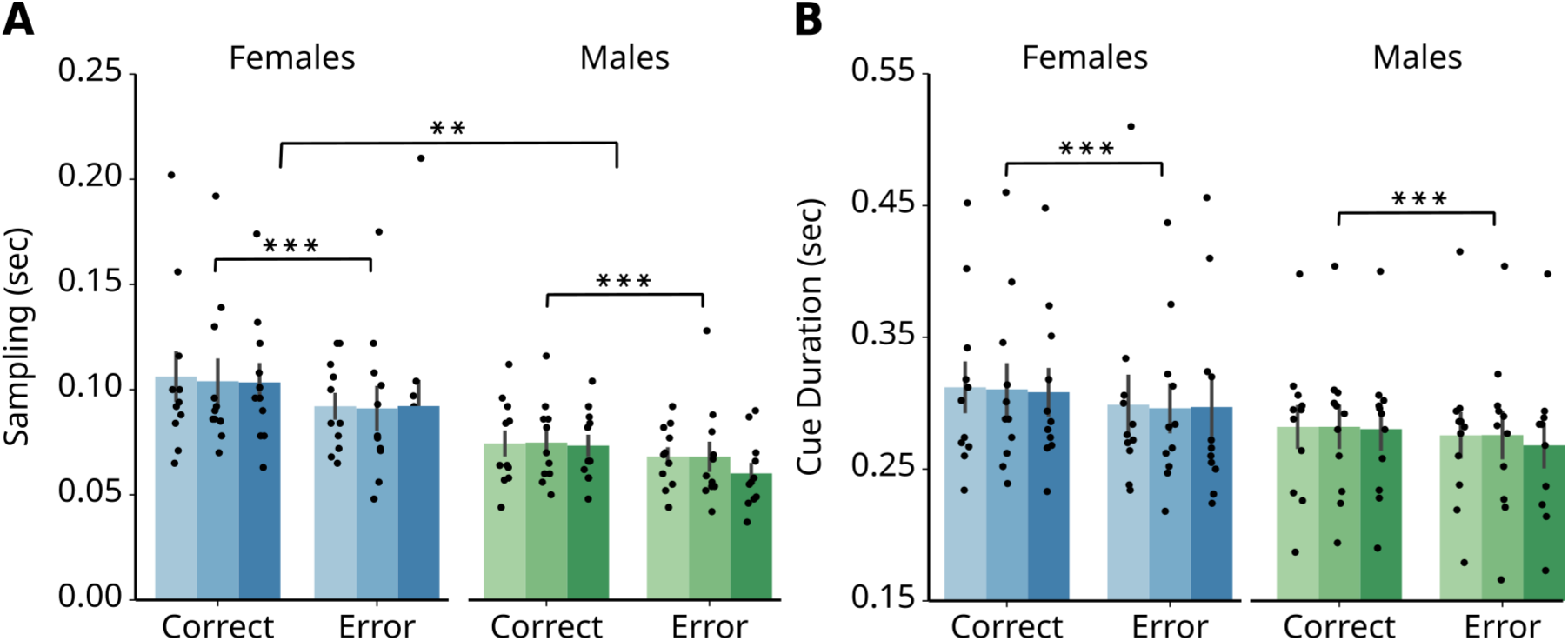
Effects of sex on active sampling. Females consistently engaged in longer active sampling than males, with all animals shortening their sampling time before making an error. **A.** Median sampling times for the main cohort of 22 animals. "Cue duration" is defined as the total time spent sampling plus the animal’s minimum response time. Females sampled for longer periods than males. Sampling times were shorter on error trials. **B.** The median cue durations did not show a significant sex difference. However, cue durations were still shorter on trials where animals made errors. Significance is denoted by asterisks here and throughout the figures: *** = p<0.001, ** = p<0.01, * = p<0.05.

As in the first cohort, sampling time did not covary with cue luminance (Figure 2B). Both sampling time (p=1.002e-05, F(1,20)=34.257) and, as a result, cue duration (p=1.816e-05, F(1,20)=31.202) were longer when animals made correct responses. Choice times, in contrast to sampling and cue duration, were longer when animals made errors (p=7.833e-08, F(1,20)=67.360). These findings replicate those of the first cohort, and further add support to our proposal of an ecological framework for decision making in freely moving animals.

All rats demonstrated an effect of cue luminance on detection rates (p=1.729e-15, F(2,40)=89.429), and there were no differences between males and females in their detection rates (p=0.385, F(1,20)=0.786). Pairwise differences revealed that detection increased with luminance, and that each level of luminance differed from one another: low- versus mid-luminance (p<.0001, t(40)=-8.682), mid- versus high-luminance (p=0.0002, t(40)=-4.469), and low- versus high-luminance (p<.0001, t(40)=-13.151) (Figure 3A). This was a key observation for interpreting the DDM results reported below.

**Figure 3.**
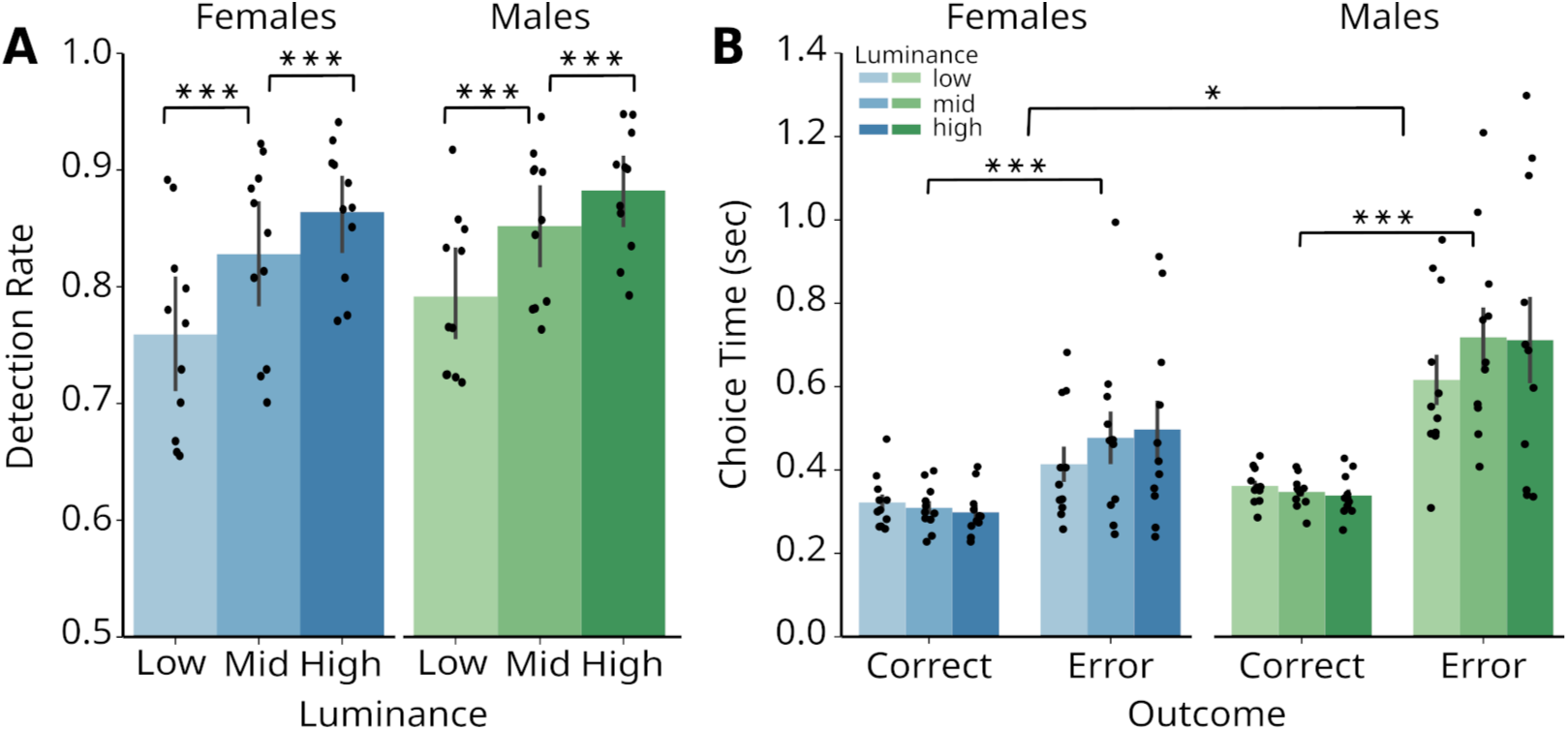
Effects of cue luminance and sex on performance. While task performance improved with higher cue luminance, cue luminance did not affect choice times. **A.** Detection rates for all animals increased as a function of cue luminance. **B.** Median choice times were not significantly affected by cue luminance. However, choice times were longer on error trials compared to correct trials, and males exhibited longer choice times than females.

A sex difference was found in choice behavior of the second cohort, as males had longer choice times (mean of 442 ms) compared to females (mean of 359 ms) (p=1.149e-02, F(1,20)=7.743) (Figure 3B). This was a difference of 83 ms on average, about a 10% difference in choice latency, which might account for a minor difference by sex in the DDM results reported below.

All animals retrieved rewards more slowly when they made errors (p=2.227e-05, F(1,20)=30.196), and there was no difference by sex (p=0.951, F(1,20)=0.004). Inter-trial intervals were similarly longer when animals made errors (p=2.530e-05, F(1,20)=29.576), meaning that animals took longer to initiate trials that ended up being errors. There was no difference in inter-trial intervals based on sex (p=0.599, F(1,20)=0.285).

### Effects of cues and sex on decision making

The results described replicate the finding of a sex difference in rats performing a value-based decision-making task (Palmer et al., 2024) and demonstrate a dissociation between the effects of cue luminance and trial outcome on detection rates and reaction times. The sex difference was largely diminished by the use of the temporally extended cues, and the main effect of trial outcome on choice time remained following that modification of the task used for the first cohort of rats.

To evaluate how these behavioral differences influenced the decision process, we performed drift diffusion modeling using the HSSM package. Our models revealed no significant sex-related differences in the parameters of the DDMs (Figure 4). We fit separate models to the data from females and males using a hierarchical mixed-effects model, such as for drift rate (v): v ∼ 0 + cue + (1|participant_id). This approach estimated parameters for each level of cue luminance, assuming a y-intercept of 0 and accounting for participant-specific random effects. We also fit models with sex as a regressor and obtained similar results (not shown). The mean estimates for the models from females and males overlapped, based on the Bayesian Highest Density Intervals (HDIs). Minor differences were observed in estimates of threshold, starting point bias, and non-decision time, which were slightly higher for males. These effects might reflect the slower choice latencies in males described above.

**Figure 4.**
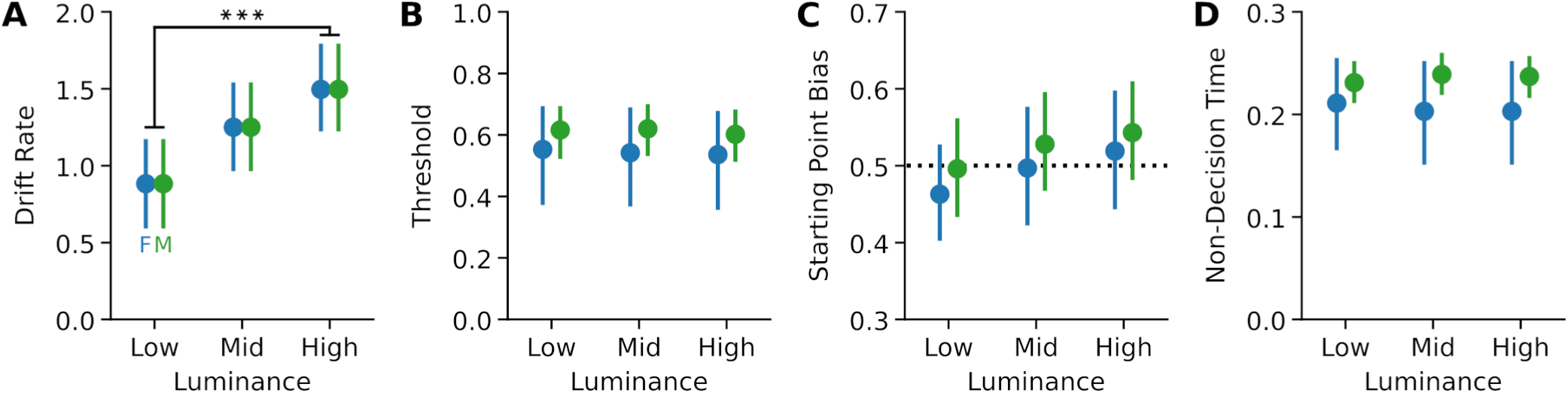
Effects of cue luminance and sex on decision making. Luminance primarily influenced the drift rate and there were no major sex differences in the DDM parameters. **A.** Drift Rate varied with the number of LEDs presented, increasing with more stimulus information. The Bayesian highest density intervals (HDIs) for drift rate estimates on trials with one LED and 16 LEDs did not overlap, indicating a significant difference. No significant differences were observed between female (blue) and male (green) rats. **B.** Threshold did not vary with cue luminance and showed no major differences between sexes. **C.** Starting Point Bias was equivalent by cue luminance and sex. **D.** Non-Decision Time (NDT) did not vary with cue luminance. The Bayesian HDIs overlapped by sex, although NDT in the models for males were longer than in the models for females.

In contrast to the lack of sex effects, the models revealed a strong effect of cue luminance on drift rate (Figure 4A), which increased with the number of LEDs presented to the rats, but not on other DDM parameters (Figure 4B-D). The lower bounds of the highest density intervals (HDIs) for drift rate estimates for the high luminance (16 LED) cue did not overlap with the upper bounds for the low luminance (1 LED) cue, with the estimates for the mid luminance (4 LED) cue falling in between. This result suggests that increasing cue luminance enhances drift rate, which may account for the primary behavioral effect of luminance on detection rates.

### Effects of cues and sampling times on detection rates

A key idea motivating this study was that in studies using freely moving animals the actions that create access to the task stimuli are part of the decision process. The time spent sampling the cues was under the voluntary control of the rats, and varied with the success of the rats in detecting the cues. To assess whether sampling time was predictive of performance, we used Bayesian Bernoulli regression in Python’s Bambi package (Capretto et al., 2022) to model the rats’ binary choices on a trial by trial basis. We fit separate models using cue luminance (1, 4, or 16 LEDs) and sampling times (from center port entrance to exit) as predictors. Models were made using data from all rats and also separately for data from the females and males. Parameter estimates did not differ across these models (and not by sex), and we report results from the total dataset below. As the behavioral results shown in Figure 3 suggest, predictions of detection covaried with cue luminance (Figure 5A). The regression coefficients for sampling times were significantly different from zero (Figure 5B). Longer sampling times led to higher detection rates, as indicated by the positive regression coefficient for sampling times.

**Figure 5.**
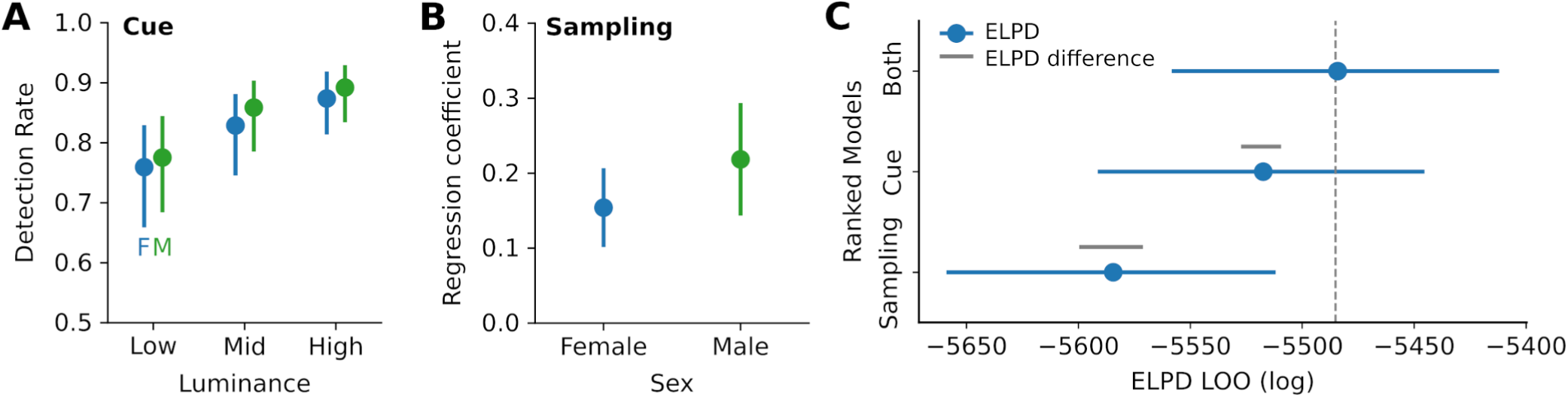
Detection rates were sensitive to both the cues and sampling time. Cue detection was most strongly predicted by cue luminance, though sampling time also contributed to predictions by the binary choice models. **A.** Models that included information about cue luminance estimated detection rates that matched those of the experimental data. Detection rates were highest for trials with the 16 LED cue compared to the 1 LED cue, as indicated by non-overlapping Bayesian highest density intervals. Results did not differ by sex. The values shown were converted from odds to probabilities for clarity. **B.** The models included sampling time as a regressor, and the regression coefficients were offset from zero for both sexes. This finding suggests that longer sampling times were associated with an increased probability of cue detection. **C.** A comparison of models based on luminance, sampling time, or both revealed that cue luminance was a stronger individual predictor of detection than sampling time. However, the best overall models were those that included both cue luminance and sampling time. Gray lines indicate the standard error of the difference in the expected log pointwise predictive density (ELPD) from Leave-One-Out (LOO) cross-validation between each model and the top-ranked model.

To better understand how cue luminance and sampling times affect detection rates, we used Bayesian model comparisons with the ArviZ package for Python (Kumar et al., 2019). We evaluated model fits using leave-one-out cross-validation based on the expected log pointwise predictive density (ELPD) statistic. The best model for predicting task stimulus detection was the one that considered both cue luminance and sampling times (Figure 5C). This model performed slightly better than a model based only on cue luminance, which in turn outperformed a model based only on sampling times.

### Effects of cues and sampling times on decision making

If sampling time was predictive of detection, it could also affect the parameters of the DDMs. To examine how sampling times affected drift rate and threshold, we used the HSSM package, as we did when evaluating sex differences. The estimates of drift rate and threshold were the same for models based on data from male and female rats. However, the models for drift rate (Figure 6A) showed a sex difference: the regression coefficient for sampling was negative and did not cross zero for female rats, whereas for males, it spanned both negative and positive values, crossing zero. In contrast, the regression coefficients for threshold were clearly offset from zero for both males and females (Figure 6B). These results suggest that sampling time had a positive effect on the decision threshold, which increased when rats sampled cues for longer times. They also revealed a sex difference, where shorter sampling times were linked to higher drift rates in females but not males.

**Figure 6.**
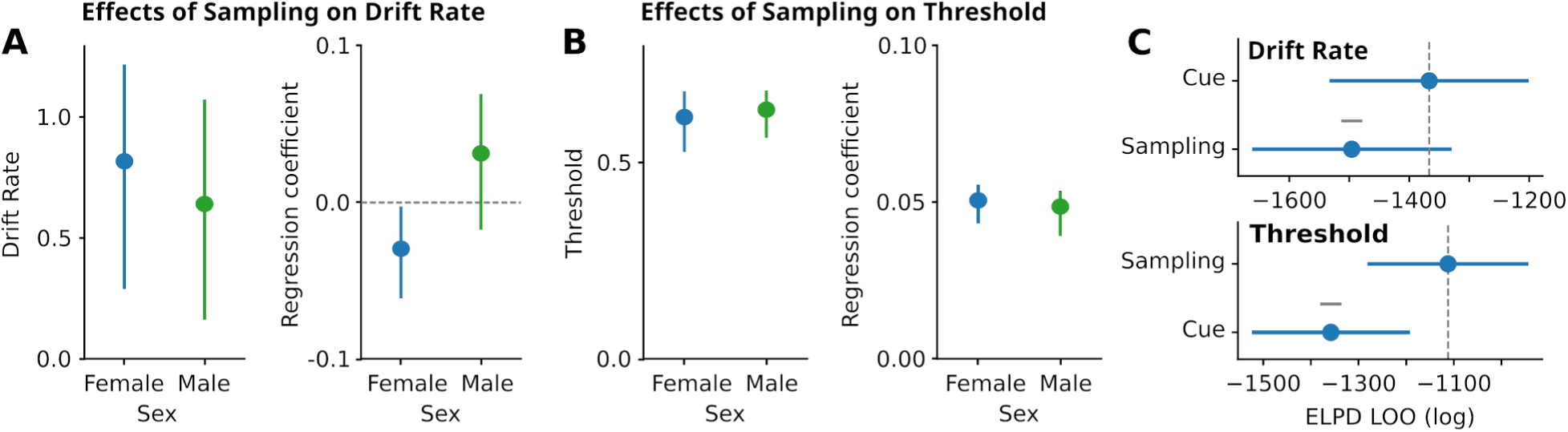
The decision threshold covaried with sampling time. Sampling time had positive influences on the decision threshold in both sexes and a unique effect on drift rates in females. **A.** Sampling time reduced drift rate in the females, but not the males. Estimates of the drift rate were similar in models based on data from the female and male rats (left). Effects of sampling time on drift rate differed by sex, with females having a negative regression coefficient for sampling time and males having a slightly positive regression coefficient (right). The Bayesian HDI for the females did not overlap zero. **B.** Sampling time was positively associated with the decision threshold in both sexes. Estimates of threshold were nearly identical for models based on females and males (left). The regression coefficients for sampling time were positive for both sexes, and the HDI did not overlap zero. **C.** Model comparisons, based on data from all rats, revealed that cue luminance was a better predictor of drift rate (top) and sampling time was a better predictor of threshold (bottom). Similar results were obtained with data only from each sex (not shown).

We compared models based on cue luminance (Figure 4) and sampling times (Figure 6) using the same leave-one-out approach described above for the Bayesian models of detection rates. We found that cue luminance provided a better model of drift rate compared to sampling times (Figure 6C, top). By contrast, sampling times provided a better model of threshold compared to cue luminance (Figure 6C, bottom). These findings collectively suggest that active sampling influences the decision process, specifically the decision threshold, as predicted by the ecological framework for decisions and not the traditional sequential framework.

## Discussion

This study examined how cue luminance and active sampling contribute to perceptual decision making in freely moving rats. We found that cue luminance selectively affected drift rate across three luminance levels, while active sampling time was associated with the decision threshold. These findings extend previous work on value-guided decisions in our lab by isolating the contribution of sensory signal strength to evidence accumulation, independent of reward value. We found clear behavioral sex differences and a sex-specific pattern in the relationship between sampling duration and drift rate. Female rats spent more time sampling the cues and estimates of their drift rates were sensitive to the time spent sampling. By contrast, males spent less time sampling and their drift rates were not detectably associated with sampling time.

### Context for and implications of the present study

This study builds upon three recent experiments in our lab that explored value-based decisions (White et al., 2024, Palmer et al., 2024, Palmer et al., 2025). These studies were motivated by Kacelnik et al. (2011), which suggested that deliberation in laboratory studies of decision making might be an artifact of the experimental design. Kacelnik’s team found that animals in nature respond to single offers of food as quickly as they do to multiple options. This observation challenges the idea that choices inherently require more deliberation. A key element in their studies was that the animals encountered single offers of stimuli more often than they were required to make choices between pairs of stimuli.

In White et al., (2024), we set out to attempt to replicate these effects using a standard neuroscience-style decision making task where rats produce cues by responding in head-entry ports and choosing between pairs of lateralized visual stimuli that varied in luminance and reward value. Only male rats were used in this study. The rats were first trained with only single offers of the stimuli, and then dual offer trials were added to the task. This allowed us to study how rats learn to make decisions for the first time in a given experimental setting. Following Kacelnik et al.’s approach, most trials (two thirds) were single offers. Contrary to Kacelnik et al.’s findings, we found that most rats slowed down when making choices between the stimuli. Using DDM models, we found that the decision threshold decreased as the rats gained experience with making choices, which we interpreted as the rats needing less information to make decisions as they gained experiences with making choices.

A follow-up study (Palmer et al., 2024) used an equal number of male and female rats. We observed sex differences in how they responded to the task. Using DDMs, we found that females were more cue sensitive, had higher drift rates, and maintained a stable decision threshold. In contrast, males showed a reduction in their decision threshold over the learning period, which we interpreted as higher choice impulsivity. A reviewer of that study noted that our design confounded reward value with stimulus salience, as brighter cues were tied to higher value rewards. To address this, we revised the task design in a third study (Palmer et al., 2025) to include a third stimulus with an intermediate brightness level. This change in the stimulus set reduced the differences in response latencies and the rats’ preference for the higher value cue. This demonstrated that cue luminance and salience play a major role in how rats make decisions in tasks that associate brightness with reward value.

Given these findings, the present study aimed to isolate how cue luminance itself affects rodent decision-making. We revised our behavioral design to present single stimuli on each trial, and using three luminance levels all linked to the same reward value. Based on findings from Palmer et al. (2024), which found that inactivating the rodent prefrontal cortex sped performance and reduced the decision threshold, we added active sampling to the task. We required rats to maintain head entries in the center port to extend the presentation of the task stimuli. As we analyzed data from the experiments, it became clear that sampling times were predictive of performance (Figure 5), but did not reflect cue luminance (Figure 2). When we then used DDMs to probe the decision process, we found it challenging to apply standard interpretations to the results. As sampling time accounted for variability in the decision threshold (Figure 6), it was challenging to interpret this effect as a purely cognitive process detached from the rats’ sensorimotor behavior. This led us to reconsider the framing of decision making models for studies in freely moving rodents.

### Active sampling as a form of information seeking

We found that active sampling time was predictive of both detection performance and the decision threshold in DDMs. Longer sampling was associated with higher thresholds, suggesting that animals that gathered more information before committing to a response were also more conservative in their decision criterion. This relationship is consistent with an active role for sampling behavior in the decision process, though the causal direction cannot be established from the current data alone. Future work manipulating sampling duration independently of cue quality will be needed to determine whether sampling duration drives threshold or reflects it.

Our results suggest that the rats actively sampled the stimuli as a form of information seeking, a modern neuroscience term with common meaning to “information pickup” in the classic ecological framework of Gibson (1966). Several recent studies have reported evidence for rats engaging in information seeking in a variety of experimental contexts (Foote et al., 2007; Yuki et al., 2017; Templer et al., 2017; Yuki et al., 2023). Our findings suggest an alternative framework for the formation of decisions by freely moving animals inspired by the work of Gibson (1966). Decisions are triggered by actions that enable perceptual processing that is coterminous with action selection. Actions that comprise the transition from sampling to choosing further reflect the ongoing decision process. Evidence for this later coregulation of decisions and actions has been reported in studies of humans (Servant et al., 2021; Weindel et al., 2021; Dendauw et al., 2024) and animals (Kane et al., 2024; Molano-Mazón et al., 2024).

We found that the sampling times were roughly half as long as the estimated Non-Decision Time (NDT) from the DDMs. The duration of NDTs across models reported in this study (150-250 ms) is longer than the latencies of visually evoked potentials in the rat visual pathways, including the visual cortex (e.g. Y. Chen et al., 2021), which typically occur in less than 75 milliseconds (about half the duration of the rats’ sampling times in the present study). If sampling behavior is purely passive, a temporal paradox emerges, as the active sampling phase would be shorter than the NDT, and information acquisition must occur before the DDM’s formal accumulation begins, which would fall within the NDT window. We propose that sampling time represents a period of active perception that is part of the decision process that indicates a transition from evidence acquisition to evidence accumulation. By this view, the DDM’s drift-rate parameter does not reflect objective stimulus strength alone, but may also reflect the efficiency of active information gathering that provides evidence for the decision process.

The validity of this framework could be tested using neural recordings in brain regions necessary for perceptual decisions. There should be changes in neural firing rates or other types of neural signatures, such as phase resets of local field potentials, around the time when animals terminate sampling and transition to action selection. Variability in neural activity at this time should correlate with variability in response times and detection accuracy in brain regions that are crucial for perceptual decisions and should be sensitive to perturbations of processing, such as drugs or optogenetic manipulations, that lead to performance deficits in decision making. We are not aware of any published studies reporting neural recordings or perturbations that demonstrate these processes specific to the moment when animals transition from sampling to choice behavior.

### Sex differences in decision making by rodents

Recent studies in animals and humans have established sex differences in learning, memory, and decision-making. In rodent studies, females show higher rates of conditioned responding and faster latencies to collect rewards in Pavlovian settings (Lefner et al., 2022), learn the values of task stimuli (C.S. Chen et al., 2021) and rules for action selection more quickly (Glewwe et al., 2026) in choice tasks, and show context dependency of episodic memory (Le et al., 2024). Female and male rats differ in decision making as well. Female rats are more risk averse (Orsini et al., 2016), more exploitative in probabilistic learning (C.S. Chen et al., 2021), less sensitive to motivational biases over instrumental actions (Degni et al., 2024), and slower to initiate trials following negative outcomes (Cox et al., 2023). These behavioral differences are linked to specific neurobiological mechanisms, such as reduced dopamine release during Pavlovian learning in females compared to males (Lefner et al., 2022), differences in functional connectivity within the striatum, hippocampus, and prefrontal cortex (Yagi et al., 2022), and differential sensitivity to noradrenergic modulation for maintaining task engagement (Rodberg et al., 2023). Collectively, these studies highlight that sex is a significant biological factor affecting the computations, neurochemistry, and circuit activity that drive complex decision-making processes.

A recent study from our lab (Palmer et al., 2024) discovered a sex difference in rats performing a value-based decision-making task. Female rats showed greater motivation for liquid sucrose rewards, as seen in the time they took to retrieve rewards. This is consistent with previous findings that female rats prefer sweet fluids more than males (Sclafani et al., 1987; Reichelt et al., 2016), are less sensitive to low-value sucrose rewards (Curtis et al., 2004), and may be less responsive to rewards in general compared to male rats (Orsini and Setlow, 2017). In our previous study (Palmer et al., 2024), females were faster to retrieve rewards than males during the early stage of training, when the rats learned the reward values of the task stimuli. Although this difference did not persist through the period of choice learning, it could indicate a motivational difference that drove some or all of the effects described in the present study.

A second notable sex difference was in the time the rats took to sample the task stimuli. Male rats sampled the cues for less time than female rats (Figure 2). This may have led to higher estimates of three parameters of the DDMs for the male rats (Figure 4) and the negative relationship between sampling time and drift rate found only in the female rats (Figure 6). These findings, along with reductions in the decision threshold of male rats during choice learning in the value-guided task (Palmer et al., 2024), suggest a sex difference in choice impulsivity (Hamilton et al., 2015; Cho et al., 2018). Viewed in the context of recent proposals that decision and movement vigor are normally co-regulated but can be flexibly decoupled according to task demands (Thura et al., 2025), our findings suggest that sex differences in active sampling may reflect broader differences in how information gathering, decision commitment, and action vigor are coordinated during choice.

## Contributions

Jensen Palmer, Kevin Chavez Lopez, and Mark Laubach designed the experiments. Jensen Palmer and Kevin Chavez Lopez performed the experiments. Jensen Palmer analyzed the behavioral data and contributed to the drift diffusion modeling. Mark Laubach performed the Bayesian choice and drift diffusion analyses. Jensen Palmer, Kevin Chavez Lopez, and Mark Laubach wrote the manuscript.

## Acknowledgments

We thank Yogita Chudasama, David Kearns, and Fany Messanvi for helpful comments on the manuscript.

